# Causes for and consequences of winning intergroup encounters in colobus monkeys (*Colobus vellerosus*)

**DOI:** 10.1101/2023.05.18.541271

**Authors:** Eva C. Wikberg, Sara Lucci, Emily Glotfelty, Fernando Campos, Pascale Sicotte

**Author notes:** Corresponding author, Address: Department of Anthropology, MH 4.03.38, UTSA, One UTSA Circle, San Antonio, TX 78249, USA.

## Abstract

The outcome of an intergroup encounter depends on the relative competitive abilities of the participating groups and the value of the resource for which they compete. However, few studies have been able to assess the consequences of winning intergroup encounters. We used behavioral and demographic data from 94 adult and subadult individuals in 8 groups of *Colobus vellerosus* at Boabeng-Fiema, Ghana to investigate the factors that predict winning intergroup encounters, and whether intergroup encounter success determines access to food and female reproductive output. In support of the hypothesis that groups with high-quality males would be more likely to win encounters, winning the encounter was best predicted by rates of displays by the alpha male. Food trees that were contested during intergroup encounters were more likely to have young leaves or other higher quality food items and to be important food species. Feeding was more likely to occur during and after the intergroup encounter if the focal animal’s group had won the encounter. The percentage of encounters won was correlated with the group’s dominance rank but not with home range size or the immature-to-female ratio. In populations such as the Boabeng-Fiema colobus in which male quality seems to be associated with winning intergroup encounters and gaining immediate access to food, one of the drivers for female transfer between groups may be differences in alpha male quality between groups and across time.

## Introduction

Intergroup encounters are characterized by aggressive behaviors over contested resources in a wide range of taxa (Boydston et al., 2001; Bruintjes et al., 2016; Cheney, 1987; Dyble et al., 2019; Hale et al., 2003; Holldobler, 1981; Majolo et al., 2020; Radford, 2008). The outcome of an intergroup encounter may be determined by asymmetries between the opposing groups’ fighting abilities or asymmetries associated with payoffs from winning (Maynard Smith & Parker, 1976; Parker, 1974), and can in turn have important implications for male and female reproductive success (Cheney & Seyfarth, 1987; Harris, 2006b; Lemoine, Boesch, et al., 2020; Lemoine, Preis, et al., 2020; Mosser & Packer, 2009).

Relative competitive ability may be determined by differences in group size, as groups with a numerical advantage are more likely to win aggressive encounters in a wide range of species (Adams, 1990; Black & Owen, 1989; Carlson, 1986; Cassidy et al., 2015; Crofoot et al., 2008; Dyble et al., 2019; Kitchen, 2004; Majolo et al., 2020; Mosser & Packer, 2009; Roth & Cords, 2016; Spong & Creel, 2004). However, group size may be a poor predictor of how many individuals participate in intergroup aggression, because in larger groups a growing collective action problem may dissuade individuals from participating (Olson, 1965; Reeve & Hoelldobler, 2007; Sweeney, 1973). Thus, the outcome of the encounter may be better predicted by the number of individuals that show intergroup aggression. The advantage of having a large group size or a more individuals showing intergroup aggression may also be moderated by the quality of the participants. For example, in wolves (*Canis lupus*) a larger number of adult males and old pack members increased the likelihood of smaller packs winning intergroup encounters against larger packs (Cassidy et al., 2015). In contrast, groups with fewer but larger adult males were more likely to win intergroup encounters in guerezas (*Colobus guereza*) (Harris, 2010). The characteristics that determine individual quality, such as age or body size, are likely to vary between study populations depending on who the main participants are in the intergroup encounters.

The outcome of an intergroup encounter may also be influenced by payoff asymmetries in which the contested resource is more valuable to one of the groups, providing a stronger incentive for the members of that group to defend the resource (Maynard Smith & Parker, 1976; Parker, 1974). This may be the case when the intergroup encounter occurs within the core of one group’s home range but outside the core area of the other group. The location of the intergroup encounter predicts the outcome in a wide range of taxa (Crofoot & Gilby, 2012; Harris, 2006b; Koch et al., 2016; Markham et al., 2012; Roth & Cords, 2016; Spong & Creel, 2004; Van Belle & Estrada, 2008). For example, capuchin (*Cebus capucinus*) groups are more likely to win encounters closer to the center of their home range (Crofoot et al., 2008; Crofoot & Gilby, 2012).

When inter-group encounters occur regularly between adjacent groups, a pattern is likely to emerge whereas one group will consistently win encounters against another in a specific set of conditions. This is likely to lead to the establishment of a dominance relationship between the groups, and to a higher predictability of winning for the dominant group (Crofoot & Wrangham, 2010). In Robinson’s (1988) study of wedge-capped capuchins (*Cebus olivaceus*), larger groups were more likely to win intergroup encounters against smaller groups, and females in larger groups had higher average fecundity, possibly because of their greater access to fruit trees. Although group size often predicts foraging efficiency, survival, and reproductive success (Cheney & Seyfarth, 1987; Mosser & Packer, 2009; Noma et al., 1998; Robinson, 1988), relatively few studies have been able to investigate the effects of a group’s dominance rank on access to resources and reproduction because of the difficulties of obtaining large-enough sample sizes (Crofoot & Wrangham, 2010).

The goal of this study is to investigate which proximate factors may explain winning intergroup encounters, whether winning predicts immediate access to food (as a short-term benefit of winning), and whether the percentage of encounters won is correlated with a group’s dominance rank, home range size, and immature to adult female ratio (as longer-term benefits) using up to 24 months of data from eight groups of black-and-white colobus (*Colobus vellerosus*) in the Boabeng-Fiema forest in Ghana. This highly folivorous primate forages mostly on large trees that occur at low densities in the forest and that represent potential targets of between-group contest competition (Saj & Sicotte, 2007). Male and female intergroup aggression includes stiff-leg displays, chases, and contact fighting (Sicotte & MacIntosh, 2004; Teichroeb & Sicotte, 2018). Male intergroup aggression is more common than female intergroup aggression in most groups and time periods (Sicotte & MacIntosh, 2004; Teichroeb & Sicotte, 2018; Wikberg et al., 2020; Wikberg, Gonzalez, et al., 2022). Males rarely form coalitions with other males (Teichroeb et al., 2014), while joint intergroup aggression among females decreases with more frequent male intergroup participation (Wikberg et al., 2014a; Wikberg, Gonzalez, et al., 2022). We hypothesize that the outcome of the intergroup encounter will be influenced by asymmetries in competitive ability and the value of the contested resource. Therefore, we expect that a group will be more likely to win an intergroup encounter if 1) the group has a numerical advantage, 2) the group’s alpha male engages more frequently in display behaviors than the opposing group’s alpha male (which we use as an indicator of relative male quality), and 3) the intergroup encounter occurs in the focal group’s core area but not the opposing group’s core area (forming a payoff asymmetry). We also hypothesize that groups will compete over access to food trees and that access to food will be an immediate benefit of winning the encounter. We predict that contested trees will be of species that make up a large percentage of the annual diet, that contested trees will have high-quality food items (young leaves, fruits, or seeds), and that individuals of the winning group will be more likely to feed immediately after the intergroup encounter. Finally, we hypothesize that patterns of wins and losses will lead to established dominance relationships between adjacent groups that regularly interact, and that this will be associated with long-term benefits. Specifically, we predict that the percentage of encounters won will be correlated with the group’s dominance rank, home range size, and the immature to adult female ratio (as a proxy for reproductive output).

## Methods

### Study population

The population of *Colobus vellerosus* (white-thighed black-and-white colobus or ursine colobus) at Boabeng-Fiema in central Ghana (7°43’N and 1°42’W) occupies a 1.92km^2^ fragment of semi-deciduous forest (Hall & Swaine, 1981). Groups vary in size from 9 to 38 and contain one or more adult males, several adult females, and immatures (Wong & Sicotte, 2006). All males disperse while only about half of the females disperse from their natal group (Sicotte et al., 2017; Teichroeb et al., 2009, 2011; Wikberg et al., 2012). All study groups have a clear alpha male (Teichroeb & Sicotte, 2010). Males form weak affiliative relationships with other males and rarely cooperate with each other (Teichroeb et al., 2014; Wikberg et al., 2012). Females form stronger social bonds than males in most groups, and females’ social networks are influenced by the presence of young infants, female immigration status, and kinship, but not by dominance rank (Wikberg et al., 2012, 2013, 2014a, 2014b, 2015). Females sometimes form coalitions against group members or extra-group members (Wikberg et al., 2014a, 2014b).

### Data collection

There were 11 groups that ranged in this study area (SI Fig. 1), and all groups were habituated to the presence of humans at a distance of 10 meters. In most cases, the monkeys were 20-40 meters away from the observers, as the observers stay on the trails while the monkeys spend most of their time in trees that can be up to 45 meters tall. The groups consisted of 1-9 adult and subadult males and 5-11 adult and subadult females (Table 1). We collected behavioral and demographic data systematically from May-August 2006, May-August 2007, and May 2008 to June 2009, from eight study groups (Table 1) and opportunistically from the other three groups in the forest whenever we encountered them. The observation period for some study groups started later than for other groups because of differences in the timing of when the group formed following a fission (NP) or when the group was habituated and all group members identified (BO and OD) (Table 1).

**Table 1.** The number of adult and subadult males and females in the groups of white-thighed colobus ranging in our study area at Boabeng-Fiema, Ghana, study period and contact time with focal groups, the percentage of intergroup encounters won, and the group’s dominance rank in the northern (N) and/or southern (S) part of the study area.

| **Group** | **# Males** | **# Females** | **Study period** | **Contact hours** | **% Wins** | **Rank** |
| --- | --- | --- | --- | --- | --- | --- |
| **AK** | 1-3 | 5 | - | - | 26.67 | 4S |
| **BO** | 2-4 | 8-11 | 09/2008 –<br>06/2009 | 185 | 39.54 | 4N |
| **BS** | 1-4 | 5-7 | 05/2006 –<br>06/2009 | 292 | 73.08 | 1S |
| **DA** | 3-9 | 7-12 | 05/2006 –<br>06/2009 | 314 | 16.67 | 6N |
| **FA** | 7 | 2 | - | - | 37.50 | 5S |
| **NP** | 1 | 5-7 | 05/2007 –<br>06/2009 | 162 | 51.72 | 2N |
| **OD** | 1-8 | 6-11 | 06/2006 –<br>06/2009 | 173 | 36.36 | 7N |
| **RT** | 4-5 | 6-7 | 05/2006 –<br>06/2009 | 333 | 55.56 | 3S |
| **SK** | 2-3 | 8 | - | - | 15.39 | 5.5 NS (5N 6S) |
| **SP** | 1-4 | 4-6 | 05/2006 –<br>06/2009 | 284 | 85.29 | 1N |
| **WW** | 3-6 | 9-13 | 05/2006 –<br>06/2009 | 416 | 40.00 | 2.5 NS (2S 3N) |

We recorded male display behaviors *ad libitum* (Altmann, 1974) to calculate the yearly rate of male display behaviors (i.e., the number of displays divided by the total observation time for the focal group). Display behaviors of male colobus, including stiff-leg displays, jump displays, and loud calls (Oates, 1977), are highly visible, and therefore we are confident that there was no systematic bias in our ability to record these behaviors among groups and individuals. The high energetic costs of display behaviors (Saj & Sicotte, 2007; Teichroeb & Sicotte, 2009) are likely to be important constraints on their frequency given the energy-saving strategy of this colobus species (Teichroeb et al., 2003). Males vary in the frequency and intensity of display behaviors, and alpha males show more frequent and longer displays than other males (Teichroeb & Sicotte, 2010). Male vocalizations or display behaviors are correlated with male body size or condition in a wide range of animal species (Davies & Halliday, 1978; Harris, 2006a; Mappes et al., 1996; Morales et al., 2003; Reby & McComb, 2003), and we use male display behaviors in this study as a proxy for male quality (Teichroeb & Sicotte, 2010).

During intergroup encounters, which we defined as when two or more groups were within 50 meters of each other (Oates, 1977; Sicotte & MacIntosh, 2004), we recorded the location of the groups, social behaviors directed to individuals in the opposing group, and the identity of interactants *ad libitum* (Altmann, 1974). We defined aggressive intergroup participation as chasing of or contact fighting with extra-group members, or performing aggressive stiff-leg or jump displays (Oates, 1977).

We collected 10-minute focal samples (Altmann, 1974) of adult and subadult females. During focal samples, we recorded feeding continuously and we used this data to determine the number of focal follows with or without feeding within one hour of the intergroup encounter. We also recorded feeding during point samples every 2.5 minutes during the focal samples, and we used the feeding point samples to calculate which plant species made up at least 5% of their diet (Saj & Sicotte, 2007).

The colobus monkeys spend most of their time feeding on large trees (Saj & Sicotte, 2007), and all large trees (i.e., with a diameter at breast height of 40 cm or more) in the study area were identified and mapped. Once a month, phenology data were collected from up to five trees of each food species from each group’s home range to record whether each plant part (flowers, fruits, seed pods, and leaves) was present (Saj & Sicotte, 2007). Whenever possible, we also recorded the phenology of contested food trees during or after intergroup encounters. We noted the focal group’s location once an hour using a map with all large trees and trails marked. We used fixed kernel density estimation to create home ranges (95% isopleth) and core areas (50% isopleth) from the one-year continuous study period, using the “ad hoc” method for the kernel smoothing parameter in the R package adehabitatHR.

### Ethical note

Our research adheres to ABS/ASAB guidelines, IPS Code of Best Practices for Field Primatology, and the laws of Ghana. Data collection was approved by the Boabeng-Fiema Monkey Sanctuary’s management committee, the Ghana Wildlife Division, and the University of Calgary’s Animal Care Committee (BI 2006-28, BI 2009-25).

### Data analyses

When analyzing the outcomes of the encounters, we considered each distinct intergroup encounter as the unit of analysis. We excluded encounters that involved more than two study groups because in such cases it was difficult to discern whether the focal group lost against one or both of the groups encountered. We also omitted encounters against non-study groups because we lacked information on their ranging patterns, group composition, and alpha male quality. We created a generalized linear mixed model with a binomial distribution and logit link function to investigate the focal group’s likelihood of winning the encounter. We coded the focal group as “winning” the encounter if they displaced the opposing group and took over the location or tree. We coded the focal group as “not winning” the intergroup encounter when the focal group was chased away by the opposing group, when the focal group attempted to but failed to chase the other group away from a certain location or tree, or when there was no clear winner of the encounter. To investigate the effects of a numerical advantage, we included the following fixed effects: relative male group size (i.e., the number of subadult and adult males in the focal group minus the subadult and adult males in the opposing group), relative female group size (i.e., the number of subadult and adult females in the focal group minus the subadult and adult females in the opposing group), and the number of males and females in the focal group that showed intergroup aggression. We did not consider how many individuals in the opposing group showed intergroup aggression, because we were not always able to reliably record this information. To investigate the effect of alpha male quality, we used relative alpha male yearly display rate, calculated as the difference in display rates between the focal group’s alpha male and the opposing group’s alpha male. To examine the effect of resource value, we used the location of the intergroup encounter in relation to the opposing groups core areas: focal core (i.e., the contested resource was in the focal group’s core area only), both/no core (i.e, the resource was located in both or neither of the two groups’ core areas), or opposing group’s core (i.e, the resource was in the opposing group’s core area only). The variance inflation factors (VIF) of fixed effects ranged between 1.06 and 1.30. We used focal group identity as a random effect. We performed all our analyses in R version 4.1.0 with the packages glmmTMB (Brooks et al., 2017) and multcomp (Hothorn et al., 2014). We used the R packages DHARMa (Hartig, 2021) and performance (Lüdecke et al., 2021) to assess model fit and calculate VIF.

We used a chi-square test to investigate whether the observed numbers of contested trees that were important (i.e., made up 5% or more of a group’s annual diet) vs. not important food species differed from expected numbers. We calculated the expected numbers by multiplying the total number of contested trees by the proportion of mapped trees that were important (or not important) food species. We also used a chi-square test to investigate whether the observed numbers of contested trees with or without high-quality food items (i.e., young leaves, flowers, fruits, or seeds) differed from expected numbers. The expected numbers were calculated by multiplying the total number of contested trees with associated phenology data by the proportion of monthly phenology tree records with (or without) high-quality food items.

We used a generalized linear mixed model with a binomial distribution and logit link function to investigate whether the presence or absence of feeding on the contested food resource during focal follows within one hour after the intergroup encounter ended was predicted by the outcome of the intergroup encounter (win, draw, loss), whether the contested food resource was located within the group’s core area, and the number of focal samples collected within one hour after the intergroup encounter ended. The VIFs for the fixed effects were 1.02-1.10. We used focal group identity as a random effect.

We created an interaction matrix with wins and losses, excluding encounters with undecided outcomes (Table 2). The AK, FA, and SK groups were not study groups, which may have led to a relatively low number of observed encounters involving these groups during the study period (Table 2). However, this would not have influenced the outcomes encounters between these groups and study groups, as the non-study groups were also habituated to the presence of humans at the typical observation distance of 20-40 meters. Because groups that ranged in different parts of the forest never encountered each other, their dyadic dominance relationships could not be inferred. For example, BS group was the highest-ranking group in the southern cluster, but it was impossible to determine its rank relative to the two top ranking groups in the norther cluster. To be able to rank the groups into a hierarchy, we divided the groups into a southern and a northern cluster with both clusters containing SK and WW groups that ranged on both sides of the Boabeng road (SI Fig. 1). We excluded VI group from both clusters as it did not typically range within our study area. The number of decided intergroup encounters within each cluster ranged from 0-29 (Table 2). For groups which were included in both the southern and northern cluster, we used their mean rank in the following analysis (Table 1). We used Spearman rank correlations to investigate whether the percentage of encounters won was correlated with: 1) the group’s dominance rank: 2) home range size; and 3) the ratio of immature individuals (i.e., infants and juveniles) to adult female calculated (N = 8 study groups). We did not include subadults in our count of immatures, because we did not always know whether they resided in their natal or breeding group (Wikberg et al., 2012). Thus, the number of subadults may not represent the reproductive output of the adult females in their current group.

**Table 2.** The interaction matrix of number of intergroup encounters won and lost among the 11 *C. vellerosus* groups at Boabeng-Fiema, Ghana (May 2007-April 2009) in the a) southern part and b) northern part of the study area, with row heading showing the winning group and column heading the losing group.

| **a) Southern cluster** |  |  |  |  |  |  |
| --- | --- | --- | --- | --- | --- | --- |
|  | **BS** | **WW** | **RT** | **AK** | **FA** | **SK** |
| **BS** | * | 8 | 8 | 0 | 1 | 2 |
| **WW** | 3 | * | 3 | 0 | 0 | 7 |
| **RT** | 3 | 3 | * | 11 | 3 | 0 |
| **AK** | 0 | 0 | 3 | * | 1 | 0 |
| **FA** | 1 | 0 | 2 | 0 | * | 0 |
| **SK** | 0 | 0 | 0 | 0 | 0 | * |

| **b) Northern cluster** |  |  |  |  |  |  |  |
| --- | --- | --- | --- | --- | --- | --- | --- |
|  | **SP** | **NP** | **WW** | **BO** | **SK** | **DA** | **OD** |
| **SP** | * | 19 | 11 | 23 | 0 | 2 | 2 |
| **NP** | 3 | * | 20 | 0 | 0 | 6 | 0 |
| **WW** | 5 | 9 | * | 0 | 7 | 1 | 0 |
| **BO** | 1 | 0 | 0 | * |  | 13 | 3 |
| **SK** | 0 | 0 | 0 |  | * | 2 | 0 |
| **DA** | 0 | 0 | 0 | 2 | 1 | * | 2 |
| **OD** | 1 | 0 | 0 | 1 | 0 | 0 | * |

## Results

We observed 189 intergroup encounters with a decided outcome involving 12 different groups. The 11 groups with established home ranges within the study area (SI Fig. 1) won between 15% and 85% of their intergroup encounters with a decided outcome (Table 1, Fig. 1). In contrast, VI group that ranged mostly outside our study area on the other side of the village only encountered our study groups twice and lost both encounters.

**Figure 1.** The number of intergroup encounters *C. vellerosus* groups at Boabeng-Fiema, Ghana was involved in May 2006 to June 2009 with the outcome win, draw, or loss.

### The outcomes of intergroup encounters and immediate access to food resources

There were 81 intergroup encounters between two study groups with complete observational data. We used these 81 intergroup encounters in our model of intergroup encounter success. The likelihood of winning the intergroup encounter increased with the alpha male display rate (Fig. 2, Table 3). Additionally, the likelihood of winning an intergroup encounter tended to decrease if it occurred within the opposing group’s core area in comparison with encounters in the focal group’s core area, but the 95% confidence interval for this effect overlapped zero (Fig. 2, Table 3). Intergroup encounter success did not differ for other comparisons of the intergroup encounter’s spatial location (Fig. 2; no/both groups’ core vs. focal group’s core: Table 3; other group’s core vs. no/both groups’ core: Tukey test, coefficient estimate = 1.20, 95% CI: -3.23 – 0.83). The likelihood of winning the intergroup encounter was not predicted by relative male group size, relative female group size, or number of aggressive males and females in the focal group (Fig. 2, Table 3). The model explained 43% of the observed variation in intergroup encounter outcome (R^2^_m_ = 0.43).

**Figure 2.** The predicted relationship between the likelihood of winning an intergroup encounter and male quality, relative number of males, relative number of females, number of males and females in the focal group that were aggressively participating, and intergroup location with shaded areas or error bars representing the 95% confidence interval.

**Table 3.**
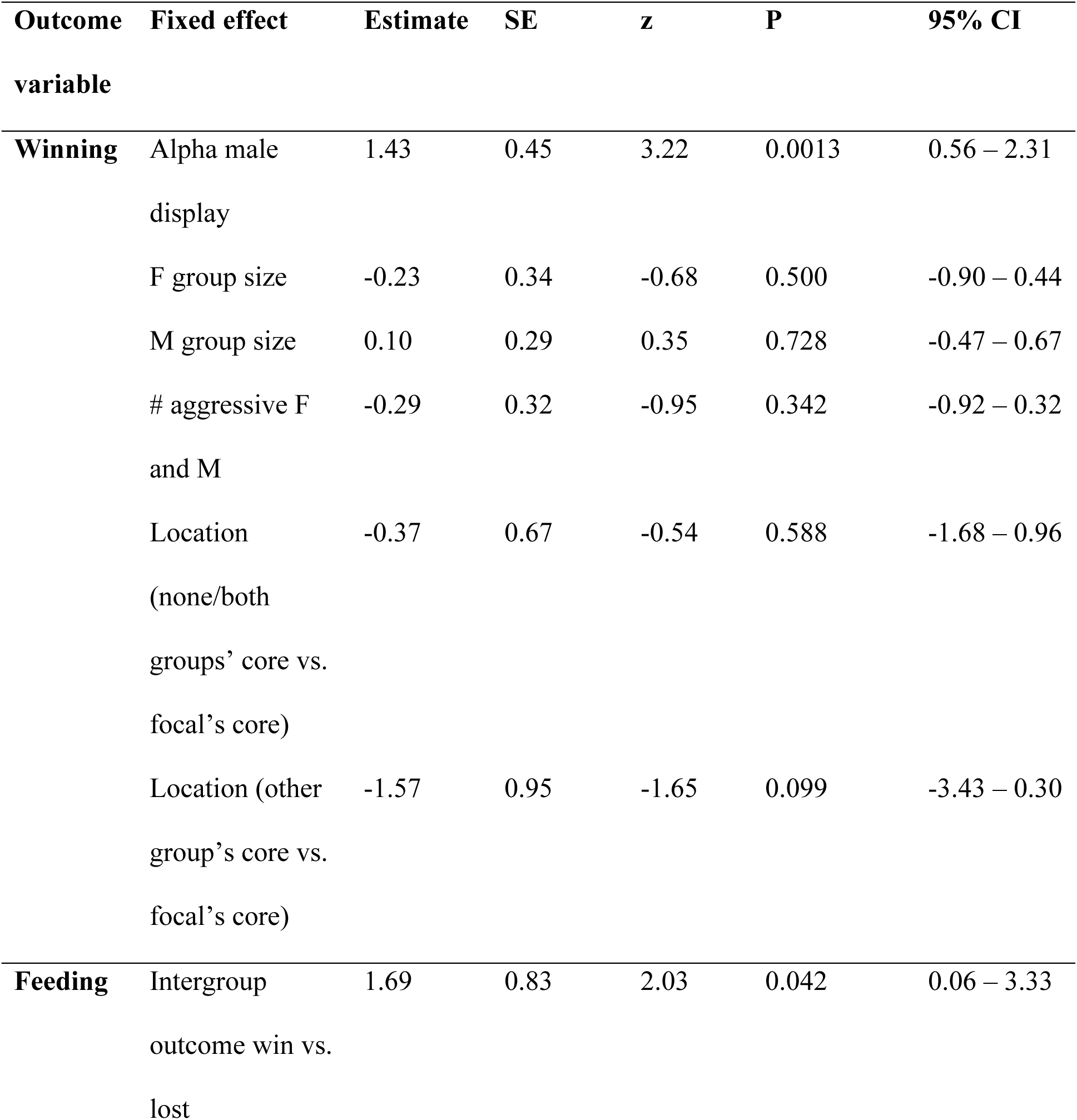
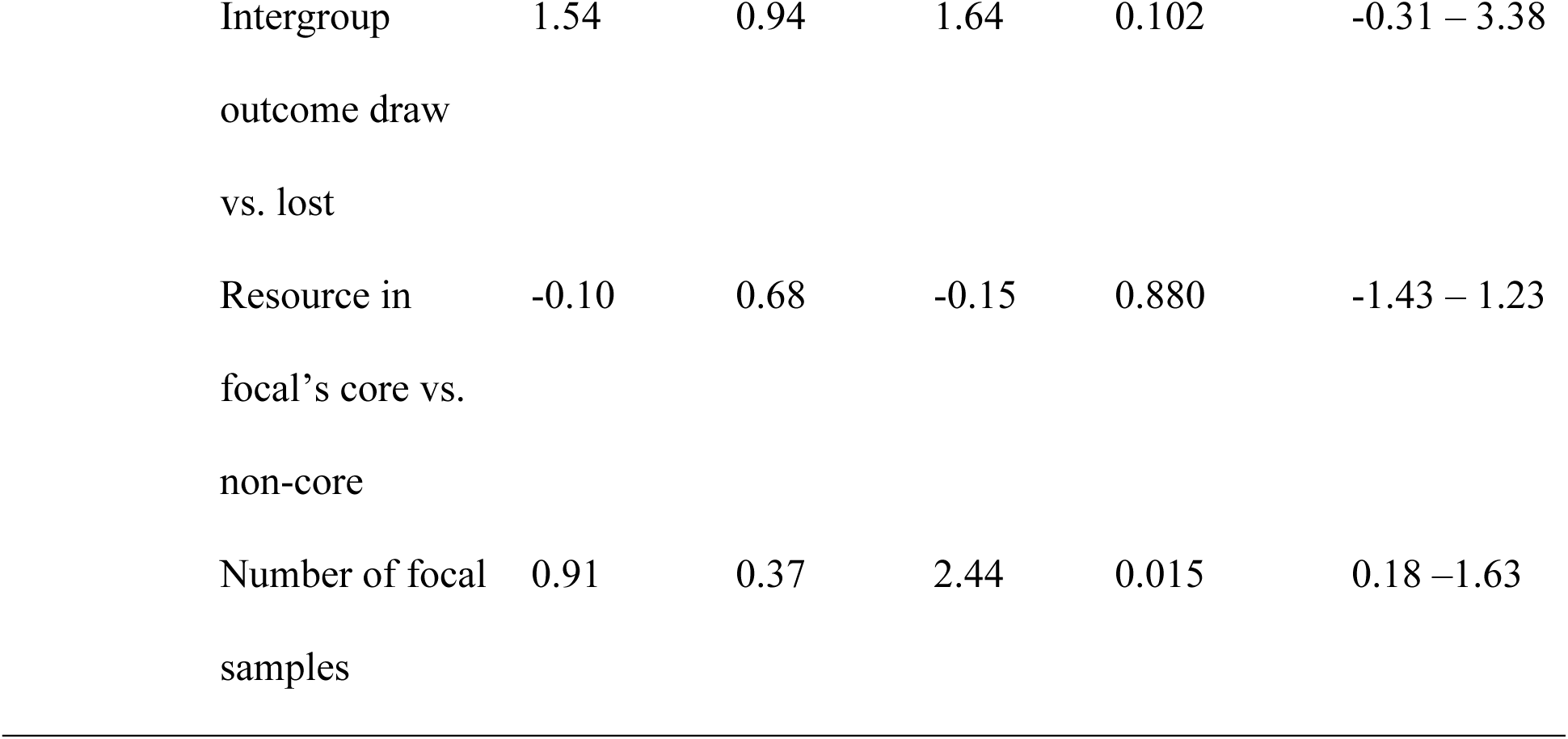
GLMMs predicting the likelihood of winning the intergroup encounter and the likelihood of feeding on the contested food resources in focal follows after the intergroup encounter in *C. vellerosus* at Boabeng-Fiema, Ghana (May 2007-April 2009).

We recorded the identity and species of the contested tree(s) in 135 intergroup encounters. The contested resource(s) included at least one tree of a species that made up 5% or more of a group’s annual diet (i.e., important food species) in 68% of these encounters (92/135). Trees that were less-important food species (<5% of annual diet) were contested in 29% of the encounters (39/135), while 3% of the encounters (4/135) did not include a contested food tree. In contrast to these patterns, only 32% of the 1,938 mapped trees were classified as an important food species. There was a significant difference in the observed versus expected number of contested trees that were important food species (chi-square = 33.85, p < 0.001). Of the 40 contested food trees for which we collected phenology data on the day of the intergroup encounter, all but one tree had high-quality food items: 19/40 trees had young leaves, 18/40 trees had young leaves and fruits/seeds, 2/40 trees had fruits/seeds; and the remaining tree only had mature leaves and no higher-quality food items. We recorded the presence of high-quality food items (i.e., young leaves, flowers, fruits, and/or seeds) in 30% of the 2,631 monthly phenology records. Accordingly, the observed numbers of contested trees with and without high-quality food items differed significantly from a null expectation (chi-square = 36.57, p < 0.001).

When analyzing whether feeding on the contested resource occurred within one hour after the end of the intergroup encounter (N = 54 intergroup encounters with focal follows), the likelihood of feeding was higher after winning an encounter compared to losing (Table 3). The likelihood of feeding did not differ after a draw compared to a loss (Table 3), or after a draw compared to a win (Tukey test, coefficient estimate: 0.16, 95% CI: -1.75 – 2.06). The likelihood of feeding was not predicted by whether the contested resource was within the group’s core area (Table 3). The model explained 25% of the observed variation in feeding on the contested resource after an intergroup encounter (R^2^_m_=0.25).

### Group dominance rank, home range size, and immature-to-female ratio

Of the 24 pairs of groups with at least one recorded intergroup encounter, in 8 pairs one group always won against the other, in 14 pairs one group won most encounters but lost at least once to the other group, and 2 pairs had tied relationships (i.e., WW won three and lost three encounters against RT; BS won one and lost one encounter against FA). The directional inconsistency index was 0.60, meaning that the higher-ranking group won over the lower-ranking group in 60% of the intergroup encounters with decided outcomes. However, 44 pairs of groups never encountered each other because several of the groups did not have neighboring home ranges (SI Fig. 1). When we constructed two separate dominance hierarchies for the southern and northern clusters of groups (Table 2), there were no intransitive relationships (i.e., no cases where group A was dominant over group B, group B was dominant over group C, but group C was dominant over group A).

The percentage of encounters won was negatively correlated with a group’s dominance rank (Spearman rank correlation r = -0.92, p < 0.001, N=11), and the highest-ranking groups in the northern and southern part of the study area won the most encounters (Table 1). The percentage of encounters won was not correlated with home range size (Spearman rank correlation r = -0.37, p = 0.362, N=8) or with the immature to adult female ratio (Spearman rank correlation r = 0.33, p = 0.420, N=8) (Fig. 3).

**Figure 3.** The relationship between the percentage of intergroup encounters each Boabeng-Fiema *C. vellerosus* group won and home range size (km^2^) and immature to adult female ratio (May 2006-June 2009). The color indicates whether the study groups ranged predominantly on the north (dark green), south (light green), or both sides of the Boabeng road (yellow).

## Discussion

In the Boabeng-Fiema colobus population, the outcomes of intergroup encounters were best predicted by alpha male display rates. Contested trees were more likely to be important food trees with high-quality food items, and winning an encounter was associated with a greater likelihood of feeding immediately after the encounter. Even though groups with high alpha male display rates were more likely to win intergroup encounters and get immediate access to food, these short-term gains did not appear to translate into long-term advantages such as occupying a larger home range or having a higher reproductive output compared to groups that were less successful in intergroup encounters. We discuss possible strategies that lower-ranking groups may have to mitigate the effects of losing intergroup encounters and having a low group dominance rank.

### The outcomes of intergroup encounters and immediate access to food

The outcome of an intergroup encounter should at least partly be determined by the relative competitive abilities of the two groups (Maynard Smith & Parker, 1976; Parker, 1974). In our study population and in other primates, residing in small groups is linked to improved cooperation during intergroup encounters (Crofoot & Gilby, 2012; Harris, 2010; Langergraber et al., 2017; Wikberg, Gonzalez, et al., 2022). However, neither relative group size nor the number of individuals showing intergroup aggression predicted the outcomes of intergroup encounters in this study, in contrast to many other studies (Adams, 1990; Black & Owen, 1989; Carlson, 1986; Cassidy et al., 2015; Crofoot et al., 2008; Dyble et al., 2019; Kitchen, 2004; Majolo et al., 2020; Mosser & Packer, 2009; Roth & Cords, 2016; Spong & Creel, 2004; Van Belle & Scarry, 2015). Instead, intergroup encounter success was predicted by display rates of the alpha male, which we used as a proxy of alpha male quality or competitive ability, because male vocalizations and other display behaviors are energetically costly and indicate male body size and condition in a wide range of species (Davies & Halliday, 1978; Harris, 2006a; Mappes et al., 1996; Morales et al., 2003; Reby & McComb, 2003), including colobus (Harris, 2006a; Teichroeb & Sicotte, 2010). It is not surprising that alpha males may influence the outcome of an intergroup encounter, because individual males, and in particular the alpha male, show intergroup aggression more frequently than do individual females (Sicotte & MacIntosh, 2004; Teichroeb et al., 2014; Teichroeb & Sicotte, 2018; Wikberg et al., 2020; Wikberg, Gonzalez, et al., 2022). Males are also the main participants in intergroup aggression in guerezas (*Colobus guereza*), and groups with fewer but larger adult males tend to win intergroup encounters against groups with more males (Harris, 2010).

The outcome of an intergroup encounter may also be affected by asymmetries in the perceived value of the contested resource. The perceived value of a resource may vary according to the location of the encounter, such that group members attribute higher value to resources located within rather than outside of the core area of the home range, and therefore are more likely to aggressively defend the resource (Crofoot et al., 2008; Crofoot & Gilby, 2012). Although our study groups tended to lose if the intergroup encounter occurred in the opposing group’s core area rather than in their own core area, the confidence interval for this effect overlapped zero, providing only weak support for this hypothesis. The location of an encounter predicts intergroup aggression and the likelihood of winning the encounter in several (Crofoot et al., 2008; Crofoot & Gilby, 2012; Furrer et al., 2011; Harris, 2006b; Koch et al., 2016; Markham et al., 2012; Roth & Cords, 2016; Spong & Creel, 2004) but not all animal populations (Cassidy et al., 2015; Dyble et al., 2019; Scarry, 2013; Spong & Creel, 2004; Yi et al., 2020). Although it is possible that these differences between studies reflect differences between populations in how intergroup encounter success is linked to the perceived value of resources, assessing perceived value can be methodologically challenging. For example, our study and several other studies did not take temporal variation in space use into account when assessing the perceived value. In our study population, specific food trees are likely more valuable when they have young leaves, although the colobus also eat mature leaves from the same trees year-round (Saj & Sicotte, 2007). For example, we have observed two groups engaging in aggressive intergroup encounters at a particular tree while it was fruiting, and yet having non-aggressive intergroup encounters at the same tree when no fruits were present (Wikberg, unpublished data). In future studies, collecting denser ranging data, and taking temporal variation in perceived value into account, will likely increase the predictive power of encounter location on the outcome. However, focusing on shorter time scales does not always improve the predictive power, as illustrated by the intensity of area use over longer time periods being a better predictor of intergroup encounter outcome than the intensity of area use over shorter time periods in a study of baboons (*Papio cynocephalus*) (Markham et al., 2012).

In our study population, contested trees had high-quality food items (i.e., young leaves, fruits, or seeds) and were of important food species (i.e., contributed to 5% or more of the annual feeding records) more often than expected based on the presence of high-quality food items in the monthly phenology surveys and the distribution of important food species in the forest. Females were also more likely to feed on the contested food tree after the intergroup encounter if their group had won the encounter. Winning the encounter therefore appears to lead to immediate access to important food resources. In other studies, the outcome of the encounter has been associated with ranging patterns that further point to immediate or short-term costs associated with losing the encounter, including increased travel and reduced use of the area after the intergroup encounter (Crofoot, 2013; Koch et al., 2016; Markham et al., 2012), and the need to shift sleeping burrow sites closer to the center of the territory (Dyble et al., 2019). Thus, individuals may benefit both from increased foraging and from reduced travel costs after winning intergroup encounters.

### Group dominance rank, home range size, and immature-to-female ratios

Groups could be ranked in a dominance hierarchy without intransitive relationships based on wins and losses, and groups that won a high percentage of intergroup encounters were high-ranking. However, we did not detect any long-term benefits from winning a high percentage of encounters and having a high group dominance rank, in contrast to the expectations of the group dominance hypothesis (Crofoot & Wrangham, 2010). The percentage of intergroup encounters won was not correlated with home range size or the immature-to-female ratio, in contrast to theoretical predictions (Isbell & Young, 2002; Koenig, 2002; Smuts et al., 1987; Sterck et al., 1997; van Schaik, 1996; Wrangham, 1980) and patterns documented in other populations (Cheney & Seyfarth, 1987; Harris, 2006b; Lemoine, Boesch, et al., 2020, 2020; Mosser & Packer, 2009; Noma et al., 1998; Robinson, 1988; Williams et al., 2004). It is possible that low-ranking groups can compensate for their low competitive ability by establishing large home ranges with little home range overlap in the secondary forest along the edges of mature forest (i.e., DA and OD in SI Fig. 1). DA and OD groups, which had the lowest dominance ranks in the northern part of the forest, have over the last four years increasingly ranged in the secondary forest along the edge of the study area as they lost parts of their home ranges close to the core forest to newly established groups, including fission products (Wikberg, unpublished data). Although the core of the study area contains more primary forest, the behavioral flexibility that this species (Wong & Sicotte, 2006) and some other African colobines exhibit (reviewed in Wikberg, Kelley, et al., 2022) may enable them to maintain high reproductive output in different habitat types. Previous studies found no difference in monthly food abundance among study groups in this forest (Teichroeb & Sicotte, 2009) or among groups in surrounding forest fragments (Wong et al., 2006), despite variation in habitat type, home range size, and degree of home range overlap. However, nutritional information is required to assess whether dietary quality differs between groups. Future studies with more longitudinal data on female reproductive output may be able to tease apart the effect of access to food from other factors that are likely to influence the production of surviving infants, such as the mother’s age (Côté & Festa-Bianchet, 2001; de Vries et al., 2016; Robbins et al., 2006; Wright et al., 2008) and male infanticide (Cheney et al., 2004; Fedigan et al., 2008; Kalbitzer et al., 2017; Manguette et al., 2019; Robbins et al., 2013; Teichroeb & Sicotte, 2008).

### Conclusions

Small group size rather than female kinship predicts female-female cooperation during intergroup encounters, which makes female dispersal compatible with cooperative home range defense (Wikberg, Gonzalez, et al., 2022). This may be particularly likely when males are frequent participants in intergroup encounters whose actions directly or indirectly defend access to food for females (Fashing, 2001; Harris, 2010; Rubenstein, 1986; Scarry, 2012; Sicotte & MacIntosh, 2004). This study further showed that female intergroup aggression is a poor predictor of the encounter’s outcome. Instead, high male display rates, which we used as a proxy for male quality, increased the likelihood of winning intergroup encounters. Winning intergroup encounters was associated with immediate access to food. Therefore, it is possible that females may transfer between groups in part as a response to differences in alpha male quality between groups and across time. By residing with a high-quality alpha male, females benefit from improved home range defense and from improved protection from infanticide.

## Acknowledgements

We thank the BFMS management committee, Ghana Wildlife Division, and the University of Calgary’s Life and Environmental Sciences Animal Care Committee for the permission to conduct this study; Robert Koranteng, Peprah Samuel, Teresa Holmes, and Fernando Campos for assistance collecting and/or organizing the ranging data; and Alberta Innovates Technology Futures (grant number 200600343), American Society of Primatologists, Animal Behavior Society, International Primatological Society, Leakey Foundation, Natural Sciences and Engineering Research Council of Canada (grant number RGPIN-203059-2006), Sweden-America Foundation, Research Services at the University of Calgary, and Wenner-Gren Foundation (grant number 8172) for funding.

## Author Contributions

ECW and PS designed the study. ECW and FAC collected, organized, and analyzed the data. EG and SL also organized the data. ECW, EG, FAC, PS, and SL wrote the manuscript.

## Data Availability Statement

Our data are stored in the PaceLab database hosted by the University of Calgary. The data used for the analyses presented here will also be available on figshare.

